# Expanded unbiased population-genomic summary statistics in pixy

**DOI:** 10.64898/2026.07.27.741083

**Authors:** Kieran Samuk, Matt Stone, Erin McAuley, Nick P. Bailey, Gina Lucas, Mikhail Plaza

## Abstract

Estimates of population-genetic summary statistics are often computed from variant-only VCFs, which commonly omit *invariant* sites (i.e. sites with only homozygous reference genotypes across all samples). However, these sites are critical for correctly estimating per site statistics such as nucleotide diversity (π) and between population divergence (d_xy_). Our software pixy addressed this issue by providing support for computing statistics directly from “all-sites” VCFs that encode invariant positions explicitly (Korunes and Samuk 2021). pixy has since been widely adopted and used in a large variety of population genetic studies. Here we present a major update to pixy, which expands the original tool in four major areas. First, we provide implementations of new estimators of Watterson’s θ and Tajima’s D that are unbiased with respect to missing data. Secondly, we provide support for arbitrary ploidy and multiallelic sites. Third, we introduce a variety of optimizations, including multicore execution and a roughly order-of-magnitude reduction in per-worker memory footprint. Finally, we have broadly modernized our code base, test suite, and community contribution pathway. We validate these new features on simulated polyploid and multiallelic datasets, as well as two empirical datasets (one diploid and one autotetraploid), and benchmark the new multicore scaling. We position pixy relative to contemporary tools and discuss shared limitations of VCF-based estimators. By integrating support for diverse ploidies, allelic complexity, and genome-wide scale in one workflow, pixy provides the population genetics community with a straightforward tool for estimates of key summary statistics that are unbiased with respect to missing data. pixy remains completely open-source (MIT-licensed) and easily installable with the conda package management system.

## Introduction

Most population-genomic analyses begin with a Variant Call Format (VCF) file, containing records of variant sites at which samples differ in allelic state from a reference genome. This is an efficient representation, but it is insufficient for estimating many population genetic summary statistics. Critically, in the absence of invariant sites, it is impossible to distinguish invariant sites (i.e. homozygous reference genotypes across all samples) from missing sites (i.e. sites where no genotypes were successfully called). This creates a variety of statistical issues. For one, summary statistics such as nucleotide diversity (π) and nucleotide divergence (d_xy_) are based on counts of pairwise differences between genotypes divided by the total number of comparisons between genotypes, including invariant sites, and are standardized by the total sequence length (Nei and Li 1979). When used to compute these statistics, a standard variants-only VCF only supplies information on genotypic differences at variant sites, divided by the total number of variant sites, which is not equivalent to π/d_xy_ as described by Nei and Li (1979). One workaround for this issue is to assume sites missing from the VCF are invariant, but this approach introduces substantial bias (Korunes and Samuk 2021). Namely, assuming that every missing *site* in the genome contains fully intact genotypes that are homozygous reference in state inflates the denominator, and assuming that missing *genotypes* at variant sites are homozygous-reference deflates the numerator (Korunes and Samuk 2021). These effects do not balance out, and their net effect is a downward bias in π and d_xy_ that scales with the amount of missing data (Korunes and Samuk 2021). We have recently shown that similar biases exist for two other related statistics, Watterson’s θ and Tajima’s D, as well (Bailey et al. 2025).

The fix for this source of bias is conceptually simple: include invariant sites in the VCF. An “all-sites VCF” encodes invariant positions alongside variant ones, so that for every window the number of missing sites and the number of missing genotypes at present sites is known exactly. With this information in hand, the denominator can be computed correctly, and estimators can be made unbiased with respect to missingness. Our software pixy (Korunes and Samuk 2021) implemented this in a command-line tool that consumes an all-sites VCF and emits per-window π, d_xy_, and F_ST_. pixy has been widely adopted, and cited by 766 works (Google Scholar, accessed July 2026). The 665 citing works indexed in OpenAlex (query in Methods) span vertebrates, invertebrates, plants, and microbes, and are concentrated in genetics, plant science, and ecology and evolution (Figure S1).

Three shifts since 2021 have motivated the expansion and updating of pixy to better serve the population genomic community. First, sample sizes have grown, and with them the fraction of sites carrying three or four segregating alleles, i.e. multiallelic sites. Sopniewski and Catullo (2024) showed empirically that the common practice of discarding tri- and tetra-allelic sites biases heterozygosity in ways that are inconsistent across species. As such, multiallelic-aware estimators are no longer optional. Second, polyploid and variable-ploidy organisms such as crop plants, allopolyploid fish, and organellar and sex-chromosome systems are increasingly the subject of population-genomic study, yet tools that natively compute π, d_xy_, F_ST_, θ, and Tajima’s D on polyploid or variable ploidy VCF files (VCFs) are sparse. genodive supports polyploids but is GUI-driven and not VCF-native (Meirmans 2020), and vcfpop targets parentage and AMOVA rather than per-window diversity (Huang et al. 2025). Finally, multicore and cloud computing are now the default, so a tool that runs single-threaded on a multi-gigabase genome is a bottleneck.

### The pixy 2.x release series

Here, we introduce a major update to pixy, which expands the tool in four areas: new summary statistics, broader data support, faster performance, and modernized engineering and community infrastructure. More than thirty distinct features and performance improvements have accumulated since the version 0.95.02 release presented in Korunes and Samuk (2021), spanning the 1.x and 2.x release series: new statistics (Hudson’s F_ST_, missingness-aware Watterson’s θ and Tajima’s D), new data support (arbitrary and variable ploidy, opt-in multiallelic sites, .csi indexes, and BED- and sites-file windowing), large performance and memory gains, and a complete engineering, testing, and packaging overhaul.

The complete pixy workflow is shown in Figure 1, with new features and workflow components tagged. A complete catalogue of all major and minor changes since the original pixy publication is given in Table S1, grouped by category (statistics, input/output, performance, dependencies, and packaging) so that headline statistical additions can be distinguished from engineering changes. Because no intervening manuscript described the 1.x development cycle, we describe all these changes together; we use ’pixy 2’ only as shorthand for the 2.x release series, and refer to the software itself as pixy throughout. All analyses reported here used the archived 2.2.3 release.

**Figure 1.**
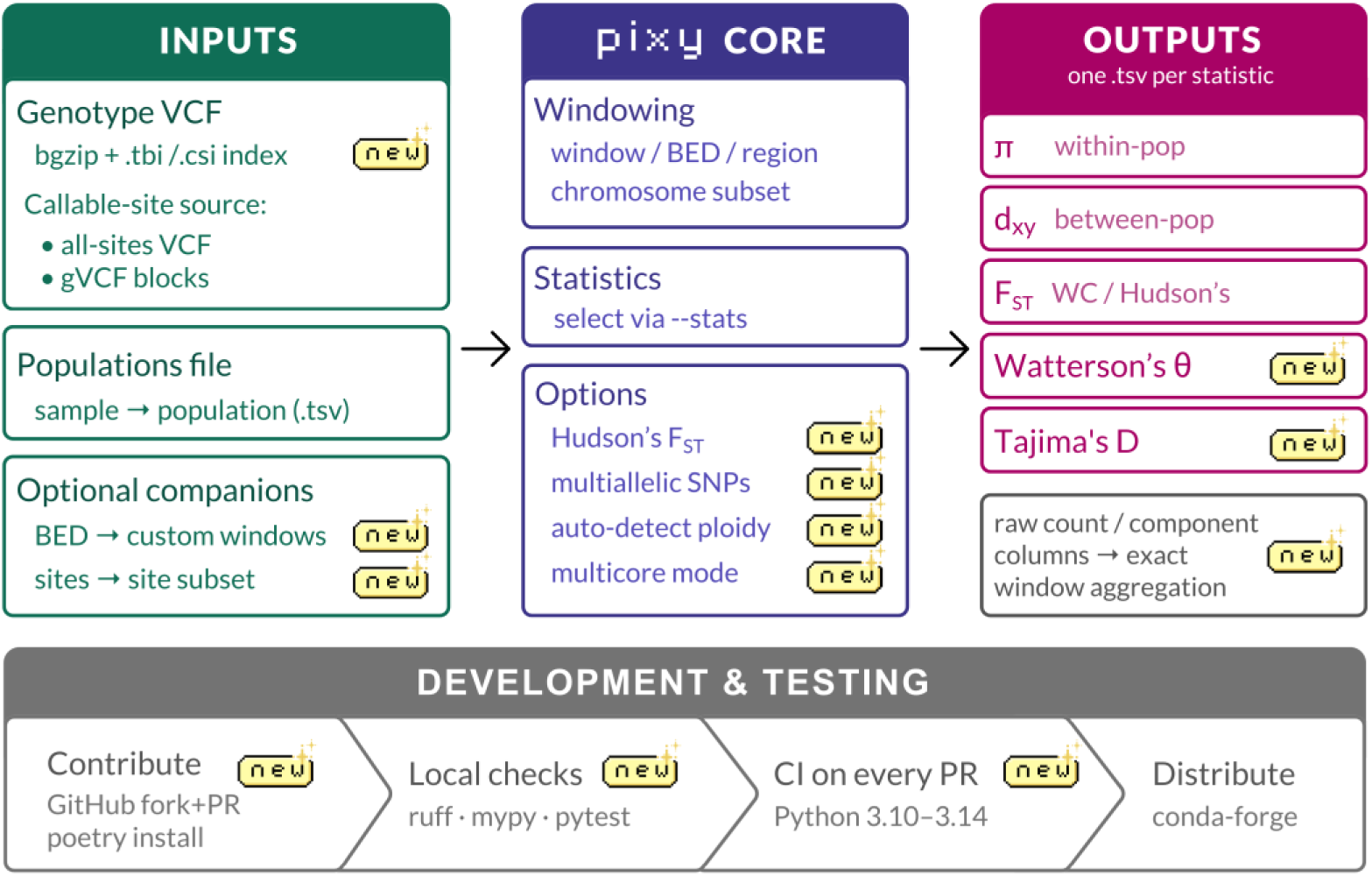
Schematic of the pixy workflow and the features added since the previous release (badged “new”). Inputs (left): a bgzipped, tabix/CSI-indexed genotype VCF, with callable sites supplied either as an all-sites VCF or as gVCF blocks; a populations file mapping samples to populations; and optional BED (custom windows) and sites (site-subset) files. Core (centre): windowing by fixed window, BED, or region with optional chromosome subsetting; statistic selection via --stats; and options including Hudson’s F_ST_, multiallelic-SNP support, automatic ploidy detection, and multicore execution. Outputs (right): one .tsv per statistic — π (within-population), d_xy_ (between-population), F_ST_ (Weir–Cockerham or Hudson’s), Watterson’s θ, and Tajima’s D — with raw-count and component columns that enable exact window aggregation. Development and testing (bottom): community contribution via GitHub fork and pull request with poetry install, local checks (ruff, mypy, pytest), continuous integration on every pull request across Python 3.10–3.14, and distribution through conda-forge.

Concretely, pixy now makes several analyses straightforward that were previously difficult or biased: estimating unbiased π, d_xy_, θ, Tajima’s D (Bailey et al. 2025), and F_ST_ in polyploid systems, and in systems whose ploidy varies among contigs (crop plants, allopolyploids, sex chromosomes, organellar contigs) from a single VCF; estimating these same quantities while retaining multiallelic SNPs; and running whole-genome and large-chromosome scans through .csi indexing, lower memory, and multicore execution, and a choice of F_ST_ estimator in one scriptable workflow. Here, we outline how each new headline feature works, provide extensive validation of each on simulated and empirical data, and discuss future work, remaining limitations, and how pixy is positioned relative to contemporary tools.

## Materials and Methods

Here, we briefly describe each new feature (Figure 1) and follow with an explanation of how each was validated against theory, simulated data, and empirical data. We point readers to Bailey et al. (2025) for the full derivations of the missingness-aware θ and Tajima’s D estimators, to Korunes and Samuk (2021) for the general π and d_xy_ framework, and to the extensive documentation at https://pixy.readthedocs.io for details of software use.

The complete source code for pixy, comprehensive Git history, and full documentation are available at https://github.com/ksamuk/pixy. The complete scripts and code used to perform the simulations and validation analyses presented here are available at https://github.com/samuk-lab/pixy-2.0.0-analysis.

### New summary statistics

Hudson’s F_ST_: pixy adds full support for the Hudson et al. (1992) estimator of F_ST_, selectable alongside the existing Weir–Cockerham estimator (Weir and Cockerham 1984). The Hudson estimator handles small and unequal sample sizes gracefully (Bhatia et al. 2013), makes native use of pixy’s existing unbiased estimates of π and d_xy_, and it now serves as the default F_ST_ estimator in pixy.

Watterson’s θ and Tajima’s D: pixy now reports Watterson’s θ (Watterson 1975) and Tajima’s D (Tajima 1989) using the missingness-aware estimators of Bailey et al. (2025), which correct for missing data in the same spirit as the original pixy’s π and d_xy_. In multiallelic mode, the segregating-site count S in both statistics is replaced by η, the parsimony-minimum number of mutations, obtained by summing (k − 1) over sites, where k is the number of alleles observed at a site (Tajima 1996). η is evaluated at the gene copies actually observed at each site so that missing data and mixed ploidy are handled correctly. On biallelic data η = S and both statistics reduce exactly to their classical forms.

Two aspects of pixy’s Tajima’s D differ from the account given in Bailey et al. (2025), as described below. The first is that the π in the numerator of D is not the general π that pixy reports for other sites. D contrasts π with Watterson’s θ on the scale of *mutation counts* (Tajima 1989), so neither term can carry a per-site denominator. The numerator is therefore the *sum of per-site pairwise diversity across sites* (Nei and Li 1979; equation 2 in Konopiński 2023), computed as in scikit-allel (Miles et al. 2021), whereas the general π that pixy reports is the sample-size-weighted average of that same quantity (Korunes and Samuk 2021; Samuk 2023). We use the unnormalised sum here because the variance term that standardizes D is itself defined for counts of mutations in a window (Tajima 1989), so dividing the numerator by per-site sample size would place it on a different scale from the quantity that standardizes it. Missing data are handled within each term by evaluating it at the level of gene copies actually observed at each site, so no further per-site weighting is required.

The second difference is a change to the formula for Tajima’s D itself. Equation 4 of Bailey et al. (2025) evaluates Tajima’s (1989) variance term separately for each class of sites sharing an observed allele count, and sums the square roots of those terms. A sum of square roots is at least as large as the square root of the corresponding sum, so this inflates the denominator of D whenever per-site allele counts are ragged, which pulls |D| toward zero and narrows its distribution. As of 2.0.0, pixy evaluates the variance term once, from the total number of mutations in a window and the mean number of alleles observed per site, rounded to an integer because Tajima’s coefficients are defined only for whole numbers of sampled alleles. The two forms agree when every site has the same observed allele count, and so diverge only in the presence of missing genotypes. We note that no released version of pixy computed the earlier form: it appeared only in 2.0.0 pre-release builds and in the version used for the analyses of Bailey et al. (2025).

### Expanded data support

Arbitrary and variable ploidy: pixy generalizes the computation of per-genotype allele counts to accept arbitrary uniform ploidies, and ploidy that varies across contigs within a dataset (numerical accuracy was evaluated for even ploidies from 2n to 8n; see Results). Ploidies may differ genome-wide (1n, 2n, 4n, etc.) or between contigs within one VCF, for example diploid autosomes coded alongside a haploid sex chromosome or organellar contig. To label each contig efficiently, pixy infers the ploidy from the first records of each contig (batched into a single index query to keep fragmented assemblies fast); it therefore assumes ploidy is constant within contigs and uniform across samples for that contig. Because pixy operates on per-site allele counts, ploidy enters the estimators only through the number of gene copies observed at each site: higher ploidy raises the number of sampled gene copies, and with it the incidence of multiallelic sites, and coarsens the granularity at which genotypes go missing, but it does not otherwise change the form of the estimators. Mixed-ploidy genotypes that vary *within* a contig, or per-sample ploidy differences on the same contig (e.g. mixed-sex genotyping of a sex chromosome in one VCF, or PAR/non-PAR transitions), can be split into separate contigs or files. Since the non-PAR regions have different effective population sizes between the sex-limited chromosomes (e.g. Y or W) versus the sex-shared chromosomes (e.g. X or Z) (Wilson Sayres 2018), they have different expectations for all the statistics calculated by pixy and therefore should be analyzed separately. Additionally, sex-biased selection and gene flow will differentially affect the expectations for these statistics (Wilson Sayres 2018).

Multiallelic sites: SNPs with more than two segregating alleles are now fully supported via the --include_multiallelic_snps option in pixy. When enabled, multiallelic SNPs are retained in the shared genotype array and so flow into every statistic: π, d_xy_, Watterson’s θ, Tajima’s D, and Hudson’s F_ST_. There is no multiallelic derivation for Weir and Cockerham’s F_ST_, but we retain the option to compute it using biallelic sites only. We note that enabling --include_multiallelic_snps changes the estimand of Hudson’s F_ST_, which then measures observed sequence dissimilarity rather than the coalescent ratio (see Results and Discussion); pairing it with --fst_biallelic retains the coalescent estimand while keeping multiallelic sites in π and d_xy_.

.csi indexes and large chromosomes: pixy now natively accepts .csi indexes in addition to .tbi, enabling analysis of chromosomes longer than 512 Mb, which are common in plants and amphibians. BED-defined windows: In addition to fixed-width windows, pixy accepts arbitrary user-defined window coordinates from a BED file. These can be used, for example, to compute π only in genic regions or at fourfold-degenerate sites, or to exclude repetitive regions of the genome from analysis.

### Performance and engineering

Multicore execution: pixy now allows for parallelized computation of statistics across windows and chromosomes. Our implementation uses Python’s standard-library multiprocessing module to dispatch jobs for each window being analyzed to worker processes and collates per-window results. Parallelism is therefore most effective when many windows are simultaneously computed across a large contig.

Shedding of zarr pre-computation: In previous versions of pixy, VCFs were converted into a database format (Zarr) prior to computing summary statistics (following the general approach of scikit-allel; Miles et al. 2021). This is efficient but introduces dependency load and one-off on-disk artifacts. pixy replaces this step with a tabix-backed reader that streams genotypes directly from the bgzipped VCF, avoiding both issues.

Modular codebase, tests, and packaging: In its original incarnation, pixy was a single large script, containing all logic and functions needed to compute summary statistics. We have completely overhauled the architecture of pixy, and decomposed the code base into modules handling parsing, statistics, I/O, and parallel-dispatch. Critically, this facilitated the development of a full test suite and continuous integration pipeline, and pixy now hosts a full pytest (Krekel et al. 2004) suite including numerical regression tests, static type checking with mypy (Lehtosalo et al. 2024), and linting with ruff (Astral Software Inc. 2024). All of these can be run locally to validate new changes, and are run automatically with every push and pull request on the pixy GitHub repository via GitHub Actions, across the supported Python versions (3.10–3.14). The project also migrated to a modern Poetry / pyproject.toml structure to standardize the build process, further easing development across different systems. These architectural changes have also greatly streamlined community contributions to pixy, by documenting environment setup, the testing workflow, and the pull-request process. These changes have facilitated expanded collaboration on pixy’s core functionality and community-submitted bug fixes.

### Simulation framework and validation datasets

To confirm that pixy introduces no changes or regressions relative to the previous release, and to validate the new headline features discussed above, we performed extensive validations on simulated data with known theoretical expectations, as well as empirical datasets with varying ploidy. All simulations were performed with our new fully validated tool vcfsim (Goulart and Samuk 2026), which wraps msprime (Kelleher et al. 2016; Baumdicker et al. 2021) to produce all-sites VCFs with precisely controlled missing data and ploidy. Simulations used vcfsim at commit da0b8ad (two commits after v1.2.0); this differs from the tagged v1.2.1 only in a population-file writing routine that this pipeline does not invoke.

### Computational configuration

All simulations and analyses were carried out on the University of California, Riverside High-Performance Computing Center (HPCC) running Rocky Linux 8.10 (Green Obsidian). Software environments were managed with conda (using the conda-forge and bioconda channels), with a separate environment defined for each analysis to pin exact versions of pixy: the legacy 0.95.01 release (code-identical to the 0.95.02 used by Korunes and Samuk (2021), the two differing only in README and documentation files) as the single-core baseline, and the latest release 2.2.3, as well as environment-specific versions of samtools, bcftools, and htslib/tabix, against a fixed Python interpreter (3.11–3.13). Analysis and simulation scripts were written in Python and R, and pipelines were orchestrated as bash job scripts submitted to the cluster’s SLURM scheduler as array jobs. Jobs were run on the HPCC’s Intel (intel) and AMD EPYC (epyc) compute partitions, with per-job resource requests scaled to the analysis (typically 1–16 CPU cores and 4–64 GB of RAM per task). The multicore benchmark (Figure 2), where absolute wall-clock and memory are compared between versions, was pinned to the Intel partition so that all of its measurements share one hardware type.

**Figure 2.**
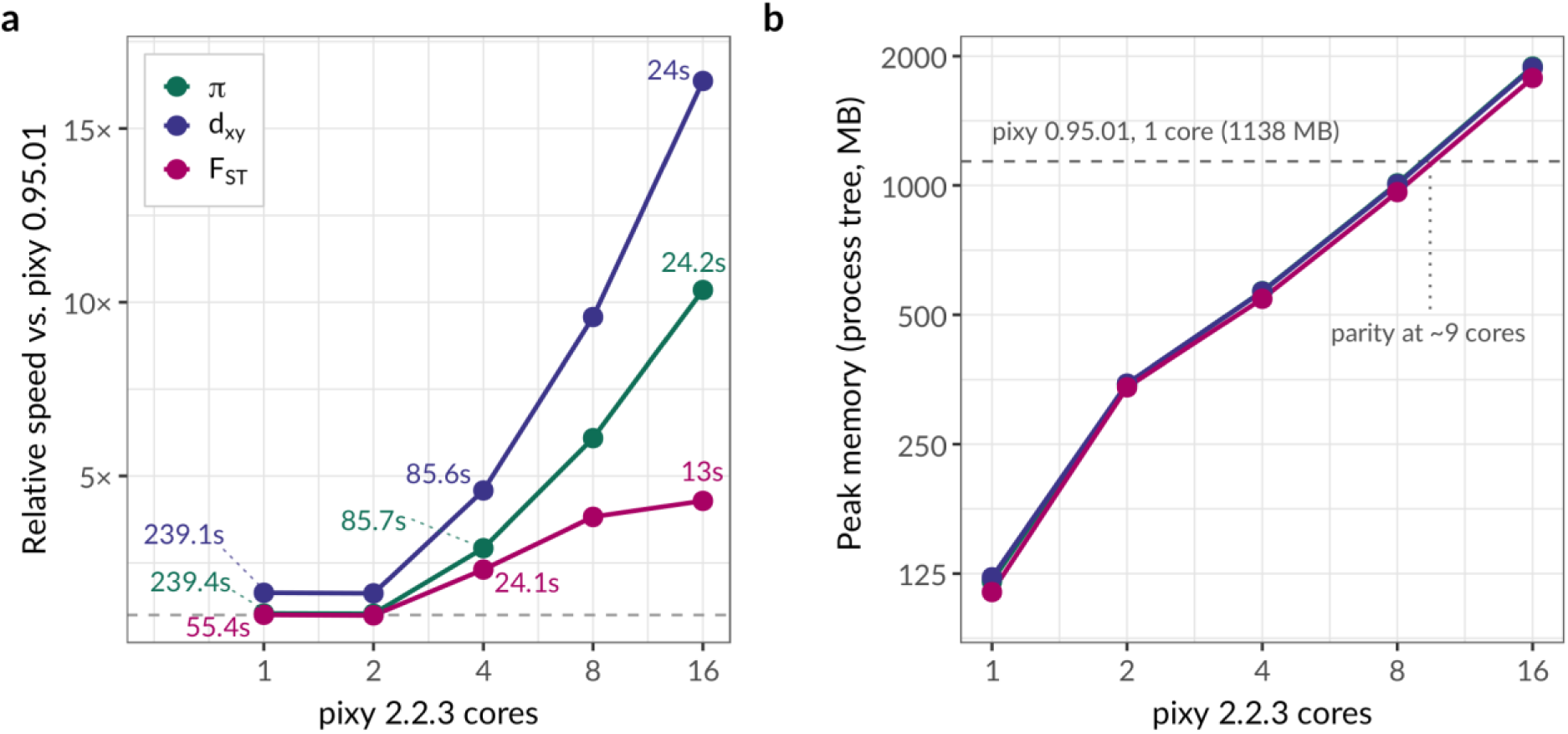
M ulticore performance and memory usage in pixy. (a) Relative speed of pixy for π, d_xy_, and F_ST_ at 1/2/4/8/16 cores vs. previous single-core release (v0.95.01), as a function of core count (log₂ x-axis; dashed line is parity). Absolute completion times in seconds are given at 1, 4, and 16 cores. (b) Peak memory usage (resident set size for the whole process tree, in megabytes) for pixy 2.2.3. The y-axis is displayed on a log2 scale. The dashed line shows peak memory usage for a single core run of pixy 0.95.01, with the dotted line showing the approximate location of parity between the two versions.

### Comparison to pixy 0.95.01

To begin, we compared the numerical outputs of pixy 0.95.01 and pixy 2.2.3 on simulated diploid data across the three statistics shared by both versions (π, d_xy_, and Weir–Cockerham’s F_ST_) at five missingness levels (0, 10, 25, 50, and 75%). To do this, we used vcfsim to simulate VCFs from populations that evolved under a shared neutral demography based on *Drosophila melanogaster* (N_e_ = 1,720,600, μ = 5.49 × 10⁻⁹, 100 kb sequences scored in 10 kb windows). π was estimated on a single panmictic population of 10 diploid samples. d_xy_ and F_ST_ were estimated using VCFs from vcfsim simulations of samples of two populations of 10 diploid samples, with a simulated divergence time of 1 million generations, giving a theoretical F_ST_ of 0.2370 under the structured coalescent (Slatkin 1991). For each simulated VCF, we produced estimates with both pixy 2.2.3 and pixy 0.95.01 in biallelic mode on identical input VCFs, and collapsed each replicate’s windows to a single per-VCF value by taking the weighted mean of the statistic across windows (using a ratio of sums approach).

### Multicore performance

Next, to assess the performance of the new multicore mode, we first simulated one hundred 10 Mb all-sites VCFs with unique seeds via vcfsim using the *Drosophila*-inspired parameters above. We then computed π, d_xy_, and F_ST_ in 25 kb windows (with --chunk_size also set to 25 kb, giving 400 chunks per VCF and 25 chunks per worker at 16 cores) for each VCF using pixy 0.95.01 (1 core), and pixy 2.2.3 at 1, 2, 4, 8, and 16 cores. For each set of estimates, we measured two metrics of computational performance: total wall time from launch to completion (in minutes), and peak resident set size (RSS, in megabytes). Because pixy 2.x dispatches one worker process per core, RSS was sampled by walking the process tree from pixy’s process identifier at fixed intervals, recording both the peak summed RSS across all live processes in the tree (the memory cost of the job as a whole) and the peak RSS of the largest single process (the per-worker footprint); the two are identical for single-process runs of the previous release.

### Statistical accuracy across ploidies

To examine potential bias and accuracy of pixy’s estimators across a range of ploidies, we simulated 100 kb all-sites VCFs at ploidies of 2n–8n (in increments of two), using population genetic parameters identical to those used in the simulations above (Ne ≈ 1.72×10⁶, μ = 5.49×10⁻⁹). To examine the effect of missing data, we repeated these simulations at five levels of missing data (0, 10, 25, 50, and 75%; both sites and genotypes, applied completely at random by vcfsim; Goulart and Samuk 2026). We simulated 10,000 replicates per ploidy at 0% missingness and 2,000 replicates per ploidy at each of the four non-zero missingness levels, giving 18,000 replicates per ploidy and 72,000 replicates per statistic across the four ploidies. We simulated VCFs for both single populations (n = 10 samples) and pairs of populations with a divergence time of one million generations as above (n = 10 samples per population). For each VCF, we used pixy to compute π, Watterson’s θ, Tajima’s D (single population files), as well as d_xy_ and F_ST_ (two-population files). Because of the finite-sites mutation model used by vcfsim/msprime and the increased number of haploid genotypes included in a polyploid VCF, multiallelic sites were common in our simulated VCFs. We thus estimated all quantities in pixy using multiallelic mode, and compared each statistic at each ploidy/missingness level to its theoretical expectation under a finite sites model (Tajima 1996). Hudson’s F_ST_ is an exception: it is reported here from pixy’s biallelic estimator, run without --include_multiallelic_snps. That is the correct estimator for the expectation it is compared against, the infinite-sites (coalescent) E[F_ST_] = 0.2370 (Slatkin 1991): biallelic filtering reduces the contribution of detectable recurrent mutation, and with it much of the homoplasy that makes F_ST_ depend on θ, so the biallelic estimator recovers the ratio of mean coalescence times (see below). The theoretical expectations for Watterson’s θ and Tajima’s D under finite sites and variable genotype missingness are non-linear, and are obtained differently for the two statistics: the Watterson reference follows the analytical finite-sites results of Tajima (1996), whereas the Tajima’s D reference is obtained by simulation under a matched coalescent null (see Supplementary Note).

To explore the effects of multiallelic sites on estimates of π, d_xy_, and F_ST_, we parameterized vcfsim to simulate a sweep over a range of per-site population-scaled mutation rates (θ = 4N_e_μ ∈ {0.005, …, 0.1}), using msprime’s default finite-sites Jukes–Cantor (JC69) mutation model. We performed this sweep at ploidies of 2n (diploid) and 8n (octoploid). We compared estimates from pixy in biallelic mode with those from multiallelic mode, and both with the theoretical expectations from the simulation under a finite sites model (see above).

Hudson’s F_ST_ required two theoretical expectations rather than one, because the biallelic and multiallelic-aware estimators target different quantities. Under an infinite-sites model, F_ST_ is a ratio of mean coalescence times and does not depend on θ (Slatkin 1991). With a divergence time of T = 10⁶ generations and N_e_ = 1,720,600 (τ = T/2N_e_ = 0.2906), and with the ancestral population α times the size of each daughter population, E[F_ST_] = (τ + (α − 1)(1 − exp(−τ))) / (τ + α); vcfsim simulates the ancestral population at twice the size of each daughter population, so α = 2 and E[F_ST_] = 0.2370. The biallelic estimator targets this value, because excluding observed multi-hit sites removes much of the homoplasy they carry. The multiallelic-aware estimator instead measures a ratio of observed sequence dissimilarity (de Jong et al. 2024), whose expectation depends on θ (Rousset 1996; Charlesworth 1998; Wilkinson-Herbots 1998) and is given by E[F_ST_] = 1 − E[π_w_]/E[d_xy_], with both terms taken as their finite-sites expectations under Jukes–Cantor (Jukes and Cantor 1969; Tajima 1996). Across the sweep this expectation falls from 0.2368 at θ = 0.005 to 0.2331 at θ = 0.100, a decline of 1.6%.

### Empirical datasets

Finally, to demonstrate the new features of pixy and further explore how including multiallelic sites affects inference, we computed summary statistics in biallelic and multiallelic mode using empirical data from two species. Each species was selected based on the existence of high-quality population-genomic data. The first was a diploid insect, the mosquito *Anopheles gambiae*, with data derived from the Ag1000G dataset (*Anopheles gambiae* 1000 Genomes Consortium 2017). For *A. gambiae*, we used data for eight randomly selected individuals from each of the BFS and KES populations, and restricted our analysis to chromosome 3R (53 Mb in length). The second was an autotetraploid plant, *Arabidopsis arenosa* (Monnahan et al. 2019). We again used eight randomly selected individuals from each of two populations (SPI and TRE), and restricted our analyses to a single large contig (LR999451.1, 25 Mb in length).

For each dataset we downloaded the raw FASTQ data, trimmed reads with fastp 0.23.4 (Chen et al. 2018), aligned to the reference genome with bwa-mem2 2.2.1 (Vasimuddin et al. 2019), and called genotypes at all sites with GATK4 (≥ 4.5) HaplotypeCaller and GenotypeGVCFs (McKenna et al. 2010; Poplin et al. 2018), with --sample-ploidy set to 2 (*A. gambiae*) or 4 (*A. arenosa*). Working from the all-sites VCF, we (i) set genotypes with depth DP < 5 to missing; (ii) removed indels and any SNP within 5 bp of an indel, since near-indel misalignment is a common source of spurious multiallelic SNPs; (iii) dropped variant sites failing the standard GATK hard-filter thresholds (QD < 2, FS > 60, MQ < 40, MQRankSum < −12.5, ReadPosRankSum < −8, SOR > 3) or displaying excess heterozygosity (ExcessHet > 54.69); (iv) masked positions of low mappability using a genmap 1.3.0 mask (Pockrandt et al. 2020; k = 100, e = 1), keeping only positions with mappability equal to 1; (v) retained sites whose total depth fell within the central 5–95% of the depth distribution; and (vi) required at least six of eight individuals to carry a called genotype in each population. The mappability, depth-band, and max-missing steps define site accessibility and were applied to variant and invariant sites alike, so that pixy’s per-window denominators remain correct. The GATK hard filters and the excess-heterozygosity filter are defined only for variant records, and were applied only to those; variant sites failing those filters were dropped altogether rather than being retained as callable.

For each species this produced one filtered all-sites VCF containing sixteen individuals (eight per population) across a single chromosome or contig, from which we computed π, d_xy_, and F_ST_ in 10 kb windows using pixy 2.2.3 in both biallelic and multiallelic mode.

### Analysis of benchmark and validation data

Statistical analysis and figure generation were performed in R version 4.4.2 (R Core Team 2024). Data manipulation and wrangling used the tidyverse suite of packages (Wickham et al. 2019), including dplyr, tibble, purrr, stringr, readr, and forcats. Figures were produced with ggplot2 (Wickham 2016), composed into multi-panel layouts with patchwork (Pedersen 2024) and ggh4x (van den Brand 2025), and formatted using scales (Wickham et al. 2025) for axes and legends and RColorBrewer (Neuwirth 2022) for palettes. Citation records for Figure S1 were obtained from the OpenAlex database (Priem et al. 2022) by querying its public REST API (https://api.openalex.org) directly with httr2 (Wickham 2026).

## Results

### Validation vs. pixy 0.95.01

The updated pixy estimates of π, d_xy_, and Weir–Cockerham’s F_ST_ reproduce the estimates of pixy 0.95.01 to within floating-point precision (Figure S2; R^2^ = 1.00 for all three statistics in every missingness panel; mean per-replicate absolute differences of 2.5×10⁻¹⁶, 2.5×10⁻¹⁶, and 2.5×10⁻¹⁵ respectively, the last a floating-point rather than integer ratio). As such, none of the architectural or mathematical changes to pixy have introduced any form of numerical regression.

### Multicore performance

pixy’s multicore and architectural improvements vastly reduce computation times across all summary statistics (Figure 2). In single-core mode, pixy 2.2.3 is as fast or faster than pixy 0.95.01 across all summary statistics: ≈ 1.05× for π, 1.64× for d_xy_, and 1.01× for F_ST_ (Figure 2a, cores = 1). In multicore mode, pixy’s median speedup over single-core mode is ≈ 1.0× at two cores, 2.8× at four, 5.8× at eight, and 9.9× at sixteen for π and d_xy_ (Figure 2a). Combining general codebase improvements with parallelism, pixy at sixteen cores is ≈10.4× (π), 16.4× (d_xy_), and 4.3× (F_ST_) faster than pixy 0.95.01 (which was single core only, Figure 2a). Computation times for π and d_xy_ scale near-linearly above two cores: fitting a power law (speedup = a·cores^b) on cores ≥ 2 gives an exponent b = 1.10 for π and 1.11 for d_xy_ (95% CIs spanning b = 1), versus 0.71 for F_ST_ (Figure 2a). The weaker scaling of F_ST_ is likely due to the computation of F_ST_ being more serial and already highly vectorized, and not requiring invariant sites.

These performance gains come in spite of two factors that work against the new version of pixy. The Zarr-backed computation of the original was itself inherently fast, so swapping in the streaming tabix reader would, on its own, have decreased performance; and pixy carries considerably more preprocessing overhead than the original (parsing ploidies, expanded validation and checks, and so on). The net speedup therefore reflects other backend optimizations that more than offset both.

Along with raw computational speed, changes in how pixy processes data have also substantially reduced peak memory usage, though the comparison depends on core count. In our test sets (10 Mb all-sites VCFs, 10 diploid samples, 25 kb windows), the new tabix-backed reader held the peak resident memory of an individual worker process at ≈105 MB regardless of core count, versus ≈1,140 MB for a single-core run of the previous release on the same data (Figure 2b), an approximately ten-fold reduction in per-worker footprint. Because each additional core runs its own worker process, however, the peak memory of the job as a whole (summed across the process tree) grows roughly linearly with core count, at ≈113 MB per added core: ≈120 MB at one core, ≈560 MB at four, and ≈1,860 MB at sixteen (Figure 2b). The total footprint of pixy 2.x therefore remains below that of the previous single-core release up to ≈9 cores, and exceeds it above that point (Figure 2b). We emphasize that this crossover reflects the previous release having no multicore mode in which to spend memory, rather than any regression in efficiency: per-worker memory is flat across core counts, and beyond ≈9 cores pixy trades memory for wall-clock time. These benchmarks reflect a single workload profile (10 Mb VCFs, ten diploid samples, one compute node); absolute times and memory will vary with genome size, sample count, ploidy, and hardware.

### Validation across ploidies

Across all combinations of ploidy and missingness, pixy’s estimates tracked their finite-sites theoretical expectations at every ploidy (2n–8n) and every missingness level (0–75%; Figure 3). Treating the 72,000 simulation replicates per statistic as draws from each estimator’s sampling distribution (Morris et al. 2019), the bias (difference between the mean estimate across each set of simulations and its theoretical expectation) was −0.013% for π, −0.001% for d_xy_, −0.008% for Watterson’s θ, −0.018% for Tajima’s D, and −0.054% for Hudson’s F_ST_, and never exceeded 0.15% for the four statistics with non-zero expectations in any single ploidy × missingness cell; for Tajima’s D, whose expectation is a small negative finite-sites floor rather than zero (−0.033 at 2n to −0.016 at 8n; see Supplementary Note), the largest single-cell deviation was 0.0028 in absolute units (Table S2). Across all of pixy’s estimators, these biases are biologically negligible in absolute terms, and well within sampling error for stochastically simulated population genetic data (Table S2). Across all five estimators, neither ploidy nor missing data had a meaningful effect on deviation from theory: although the large replicate count rendered a main effect of ploidy and its interaction with estimator statistically detectable (ploidy F_3, 359900_ = 15.40, p = 5×10⁻¹⁰; statistic × ploidy F_12, 359900_ = 2.20, p = 0.009), every such term explained ≤0.01% of the variance in deviation from theory. The missingness main effect and all its interactions were non-significant (missingness F_4, 359900_ = 0.67, p = 0.61; ploidy × missingness F_12, 359900_ = 1.71, p = 0.057; three-way F_48, 359900_ = 0.21, p = 1).

**Figure 3.**
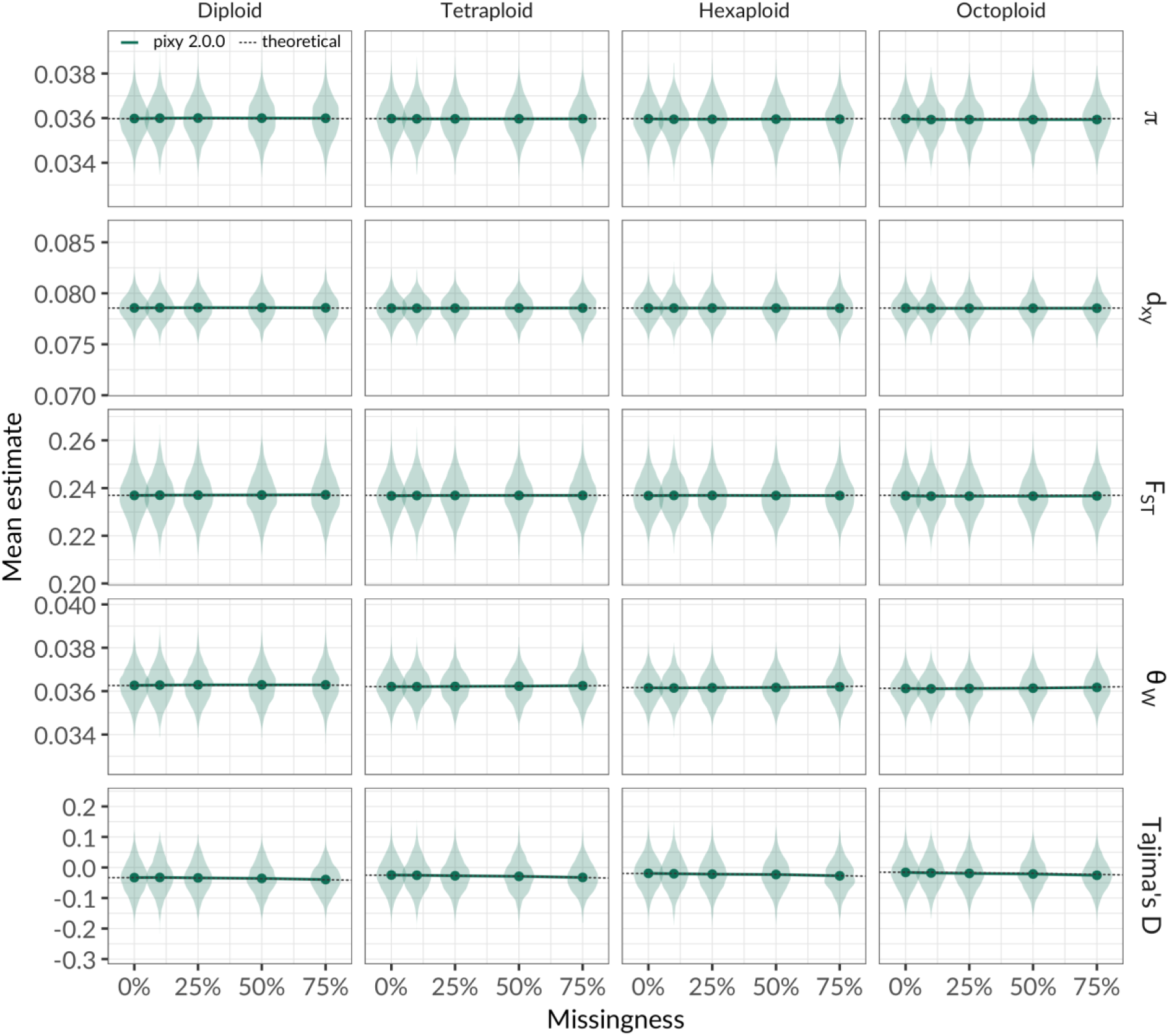
Polyploid support across four ploidy levels. The distribution (density violins) and mean (points) of pixy’s estimates of π, d_xy_, Watterson’s θ, Tajima’s D, and Hudson F_ST_ (rows, y-axis values; plotted in the order π, d_xy_, F_ST_, θ_W_, Tajima’s D) across four ploidy levels (columns) as a function of missingness fraction (0, 10, 25, 50, and 75%, x-axes). Theoretical expectations under a finite-sites model are shown for each quantity as a dotted line. For π, d_xy_, and Watterson’s θ these are the analytical Jukes–Cantor expectations of Tajima (1996), whereas Tajima’s D is instead obtained by coalescent simulation under a matched null (see Supplementary Note). π, d_xy_, Tajima’s D, and Watterson’s θ were estimated using pixy’s multiallelic-aware estimator. Hudson F_ST_ is estimated using biallelic sites, and the dashed line is the infinite-sites (coalescent) expectation E[F_ST_] = 0.2370 (Slatkin 1991) rather than a finite-sites value.

### Multiallelic-site support

The multiallelic site-aware estimators implemented in pixy eliminate biases in the estimation of π and d_xy_, particularly when the population-scaled mutation rate (θ = 4N_e_μ) exceeds 0.025 (Figure 4). Across the range of simulated values of θ (0.005–0.10) and for both ploidy levels tested, multiallelic-aware estimates of π and d_xy_ closely tracked theoretical finite-sites expectations, whereas the default biallelic-only estimates became progressively more negatively biased at higher levels of θ (Figure 4). The magnitude of the bias in the biallelic estimator also rose as a function of ploidy: for example, the biallelic bias in π ranged from −12.5% at θ = 0.1 and 2n to −17.6% at θ = 0.1 and 8n. For d_xy_, the bias was even worse, ranging from −24.8% at θ = 0.1 and 2n to −31.0% at θ = 0.1 and 8n. An analysis of variance shows that the magnitude of bias in the biallelic estimator was significantly associated with θ (F_4, 39980_ = 582.1, p < 2×10^−16^) and ploidy (F_1, 39980_ = 81.8, p < 2×10^−16^), and was significantly larger for d_xy_ than for π (F_1, 39980_ = 510.6, p < 2×10^−16^), with the θ effect itself steeper for d_xy_ (statistic × θ, F_4, 39980_ = 45.3, p < 2×10^−16^). Enabling multiallelic-aware estimation removed this bias: across every statistic, ploidy, and level of θ, the multiallelic-aware estimators of π and d_xy_ stayed within 0.8% of their theoretical expectations (pooled bias: +0.5% for π and +0.0% for d_xy_), and showed no significant relationship with θ, ploidy, or statistic (ANOVA as above, θ: F_4, 39980_ = 0.00, p = 1.00; ploidy: F_1, 39980_ = 0.23, p = 0.63; statistic: F_1, 39980_ = 2.04, p = 0.15; all interactions p ≥ 0.63).

**Figure 4.**
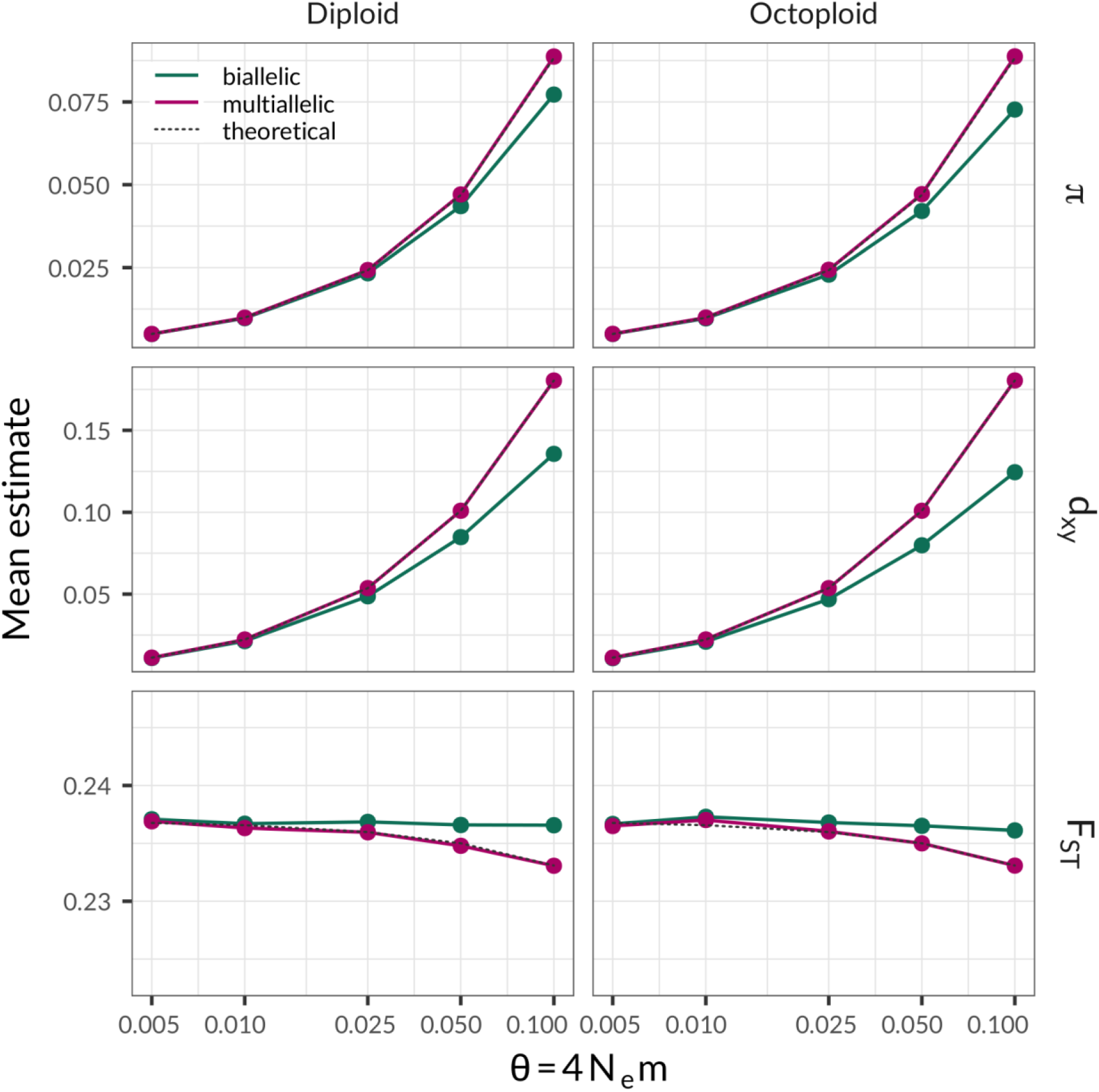
M ultiallelic-site support and H udson F_ST_ across a range of θ values. Mean estimates of π, d_xy_, and Hudson F_ST_ (rows) as a function of θ = 4Neμ, at 2n and 8n (columns), for the biallelic-only and multiallelic-aware estimators (colors). The dotted line is the theoretical expectation under the finite-sites Jukes–Cantor model used in the simulations, in every row; for F_ST_ it declines from 0.237 at θ = 0.005 to 0.233 at θ = 0.1 rather than being constant. The biallelic F_ST_ lies above that line because it recovers the infinite-sites (coalescent) F_ST_ (0.237, independent of θ) rather than the observed-dissimilarity quantity plotted for π and d_xy_; each estimator is essentially unbiased for its own expectation (see Results).

The multiallelic site-aware estimator of Hudson’s F_ST_ also reduced bias in the estimation, though by far less than for π and d_xy_ (Figure 4). Judged against the finite-sites expectation, multiallelic-aware F_ST_ closely tracked theory across the whole sweep (pooled bias −0.04%, 95% CI −0.12% to +0.04%; maximum absolute bias 0.19%), whereas the biallelic-only estimator became progressively more positively biased as a function of θ, increasing from +0.14% at θ = 0.005 and 2n to +1.49% at θ = 0.1 and 2n. An analysis of variance shows that the two estimators differ significantly in their degree of bias (F_1, 39980_ = 88.4, p < 2×10^−16^), and that the bias is significantly associated with θ (F_4, 39980_ = 21.4, p < 2×10^−16^). These effects are significant but small in absolute terms: the bias of the biallelic estimator of F_ST_ does not exceed 1.5% (even when θ is high, θ = 0.1 and 2n). This is an order of magnitude less than the biallelic deficit in π (−12.5% at θ = 0.1 and 2n) or d_xy_ (−24.8% at θ = 0.1 and 2n). Unlike π and d_xy_, the biallelic estimator’s bias in F_ST_ does not appear to be associated with ploidy (F_1, 39980_ = 0.04, p = 0.85), and there is no significant estimator × ploidy interaction (F_1, 39980_ = 0.55, p = 0.46), or ploidy × θ interaction (F_4, 39980_ = 1.75, p = 0.14).

However, this result inverts if the *coalescent* F_ST_ is used as the target estimand: compared to the infinite-sites coalescent expectation E[F_ST_] = 0.2370 (Slatkin 1991), it is the multiallelic-aware estimator that is biased, increasingly so with θ (up to −1.64% at θ = 0.1), while the biallelic estimator remains essentially unbiased across the range (maximum absolute bias of 0.35%). Which estimator is unbiased therefore depends on which of the two estimands is the target of the analysis (see Discussion).

Together, these results show that multiallelic-aware estimation in pixy removes a substantial bias in π and d_xy_, as Sopniewski and Catullo (2024) found for heterozygosity, and leaves Hudson’s F_ST_ essentially unchanged (depending on the estimand sought).

### Empirical validation on real published data

Based on the results of our simulations above, i.e. that a purely biallelic estimator tends to underestimate genetic diversity, we reasoned that applying the multiallelic-aware estimator of π to empirical data would raise these estimates toward less biased values (following Sopniewski and Catullo 2024). To ensure that any such increase reflected biological variation rather than alignment or genotyping artifacts, we computed both estimators from stringently filtered, callable-site VCFs (see Methods). As predicted, the multiallelic-aware estimator of π was higher than the biallelic-only estimator in every population we examined. The mean per-window increase was ≈8% in both populations of *A. gambiae* (BFS +8.2%, SD 4.8%, n = 5,280 windows; KES +8.0%, SD 4.7%, n = 5,279; Figure 5) and ≈9% in *A. arenosa* (SPI +9.1%, SD 8.3%, n = 2,366; TRE +9.4%, SD 15.6%, n = 2,373). Per-window d_xy_ shifted upward by a similar amount (+8.0%, SD 4.5%, n = 5,283 windows in *A. gambiae*; +9.5%, SD 26.1%, n = 2,380 in *A. arenosa*). Critically, the per-window uplift in π (multiallelic π – biallelic π) scaled with the local percentage of multiallelic sites (multiallelic SNPs as a percentage of comparable sites; Spearman ρ = 0.95 and 0.94 in BFS and KES; 0.88 and 0.90 in SPI and TRE; all p < 2×10⁻¹⁶). This mirrors the general results of Sopniewski and Catullo (2024), and, in light of the simulated results in Figure 4, indicates that the biallelic estimate of π is likely deflated in these species, and that this deflation persists under stringent site filtering.

**Figure 5.**
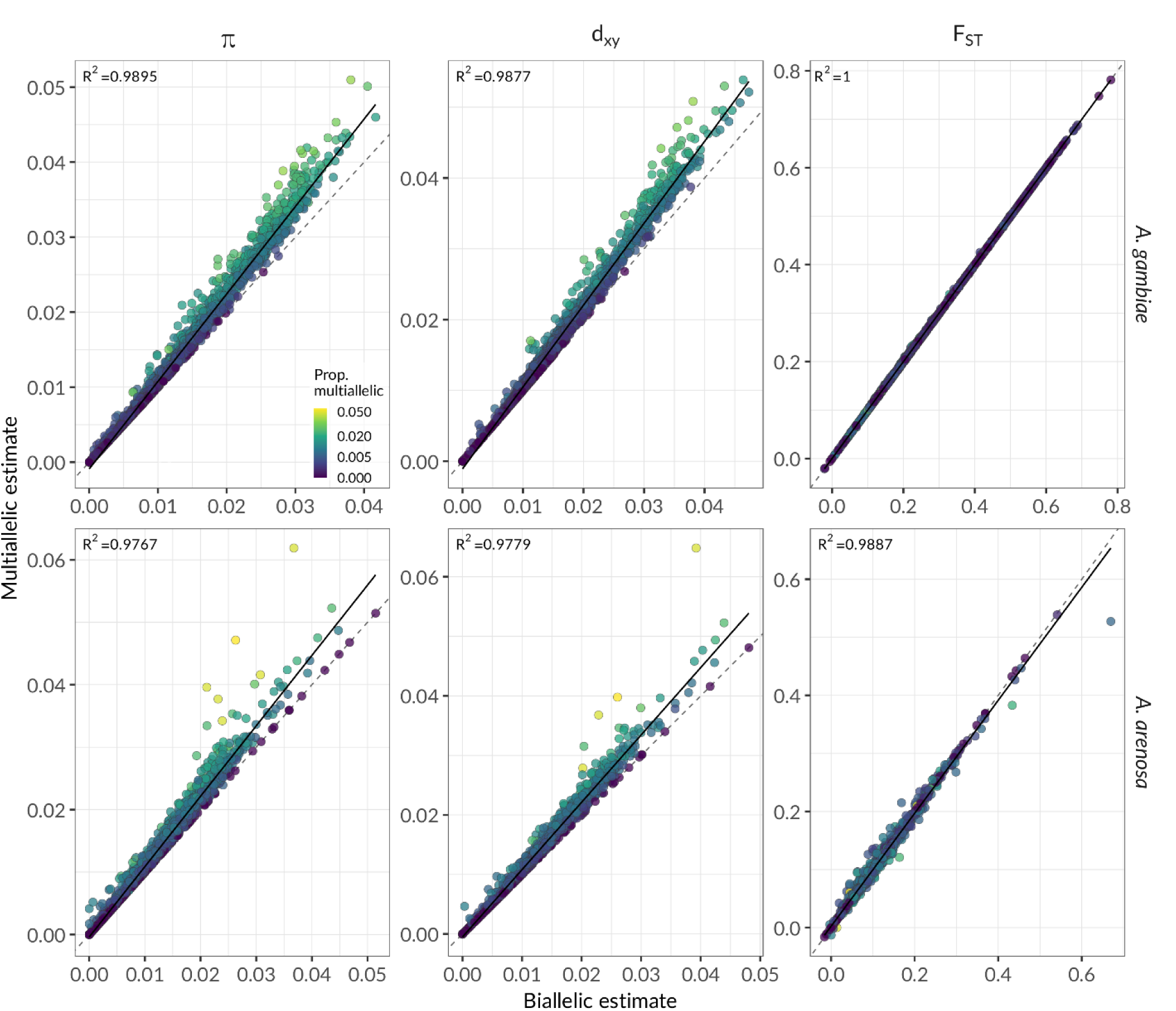
Comparison of multiallelic and biallelic estimators across two empirical datasets. pixy was run with and without --include_multiallelic_snps on GATK all-sites VCFs from *Anopheles gambiae* (2n, chromosome 3R; BFS and KES) and *Arabidopsis arenosa* (4n, LR999451.1; SPI and TRE). Per-window estimates of π, d_xy_, and F_ST_ (columns) are plotted with biallelic-only (x) against multiallelic-aware (y), one row per species. Points are coloured by the proportion of multiallelic sites in the window; the dashed grey line is the 1:1 line for both estimators and the solid black line is an ordinary least-squares fit, whose R² is given in each panel.

When comparing the multiallelic vs. biallelic estimators of Hudson’s F_ST_, per-window F_ST_ was centred on essentially no change in both species, but with far greater per-window scatter in the autotetraploid (*A. gambiae*: mean difference +2.7×10⁻⁶, SD 5.9×10⁻⁴, n = 5,283 windows; *A. arenosa*: −2.1×10⁻⁴, SD 5.8×10⁻³, n = 2,380; Figure 5). This ten-fold difference in scatter tracks the local density of multiallelic sites: in multiallelic mode the F_ST_ computation gained 0.5 additional multiallelic SNPs per window in *A. gambiae* but 12.6 in *A. arenosa*. The diploid result matches the previous finding that multiallelic-aware estimation affects π and d_xy_ while leaving Hudson’s F_ST_ essentially unchanged, and the equivalence weakens as multiallelic sites become common (as in a tetraploid).

## Discussion

Estimating population genetic summary statistics is a key part of most evolutionary and population genomic analyses. Here, we performed extensive validation of pixy’s new headline features. We showed that (1) pixy’s estimators recover their theoretical expectations across variable ploidy and missingness, (2) multiallelic-aware estimation removes a substantial bias in π and d_xy_, and (3) the rewritten architecture is both faster and substantially lighter in per-worker memory footprint than pixy 0.95.01. Below, we discuss how these results will facilitate and improve empirical practice, and how pixy sits relative to the current landscape of population genetic software.

### When multiallelic-aware estimation matters

Perhaps the most immediately useful result presented here is that ignoring multiallelic sites biases π and d_xy_ downward, and that the magnitude of this bias is primarily a function of the population-scaled mutation rate, θ. The bias also increases with the number of sampled gene copies, and hence with ploidy (see below). This is consistent with theory, and tracks the empirical results presented in Sopniewski and Catullo (2024). When θ is low, the bias introduced by ignoring multiallelic sites is low, but it can increase substantially if θ is high (up to a 31% underestimate, in our results). Enabling multiallelic-aware estimation removes this bias, to within 0.8% of theory.

Whether this matters in practice for any particular analysis will depend on the value of θ in the species under study. In both *A. gambiae* and *A. arenosa*, the multiallelic-aware estimator raised π by roughly 8–9% (Figure 5), and the uplift in individual windows tracked the density of multiallelic sites in those windows. Neither species was chosen for unusual diversity, although genetic diversity has been shown to be relatively high in both of these species, with estimates of θ ≈ 0.015 in both species (*Anopheles gambiae* 1000 Genomes Consortium 2017; Monnahan et al. 2019). This places them around the 70^th^ percentile of species, compared to species in the cross-taxa θ dataset compiled by Buffalo (2021). As such, and once again in accordance with Sopniewski and Catullo (2024), reported estimates of genetic diversity are likely deflated in studies of sufficiently diverse taxa that discard multiallelic sites. We note that analyses based on alignments or genotype likelihoods, or on any workflow that retains multiallelic information, need not share this bias.

Along with an effect of θ, the bias also grows with ploidy per se. Note that in our simulations, the haploid effective population size for all simulations was identical, so there was no confounding of ploidy and θ (although in real populations this would likely be a factor). The cause of the ploidy effect is instead sample size: under a finite-sites model, sampling more haploid genomes lengthens the genealogy and improves detection of rare alleles, so recurrent mutations at the same site are both more numerous and more likely to be observed. As such, simulated polyploid VCFs contain more multiallelic sites than diploid VCFs simulated under otherwise identical parameters. Real polyploid systems are thus the most likely to be affected by biases introduced by ignoring multiallelic sites and the least well served by existing software.

Given these results, our general recommendation is to enable multiallelic mode when estimating π, d_xy_, Watterson’s θ, and Tajima’s D in pixy. The option carries no meaningful computational cost, and produces results identical to the biallelic estimator when a dataset lacks multiallelic sites. That said, these recommendations and our results above assume multiallelic sites are actual biological phenomena, and not sequencing or alignment artifacts, which are known to introduce bias into summary statistics (Achaz 2008). As such, proper filtering, particularly of repetitive and low-accessibility sites in the genome, remains critical (Brault et al. 2026). The multiallelic forms of Watterson’s θ and Tajima’s D are finite-sites generalizations, and their interpretation differs from the standard statistics in one respect worth stating plainly. Because η counts the parsimony-minimum number of mutations rather than the number of segregating sites, η ≥ S at every site, and the resulting Tajima’s D settles on a small non-zero floor under neutrality rather than on zero (Roychoudhury and Wakeley 2010; Bhaskar, Kamm and Song 2012; see also Supplementary Note). The correction is therefore consistent rather than strictly unbiased. On biallelic data the distinction disappears, since η = S and the statistic reduces to Tajima (1989). We note that comparisons of Tajima’s D across studies should state which counting definition was used, as the two are not interchangeable at high θ.

### F_ST_ in the presence of multiallelic sites

Our F_ST_ results require more care to interpret than those for π and d_xy_, because the biallelic and multiallelic-aware estimators do not target the same quantity. Judged against the finite-sites expectation, the multiallelic-aware estimator of F_ST_ closely tracks its finite-sites expectation across all the parameter combinations we tested (pooled bias −0.04%) while the biallelic estimator drifts mildly upward as a function of θ, peaking at +1.49% at θ = 0.1. However, when both estimators are judged against the infinite-sites coalescent expectation of E[F_ST_] = 0.2370 (Slatkin 1991), the result inverts exactly: the biallelic estimator is essentially unbiased (maximum absolute bias 0.35%) and the multiallelic-aware estimator reads as slightly biased by −1.64% at θ = 0.1.

This is expected behavior of these estimators, and our results support that our implementation of both estimators is faithful and unbiased, but for different estimands. Biallelic filtering reduces the homoplasy that makes F_ST_ depend on θ (Kimura 1969; Cutter et al. 2013), so the biallelic estimator of F_ST_ recovers the coalescent ratio of mean coalescence times, whereas the multiallelic-aware estimator consistently measures the ratio of observed sequence dissimilarity (de Jong et al. 2024), which naturally changes as a function of θ under a finite-sites model (Rousset 1996; Charlesworth 1998; Wilkinson-Herbots 1998).

In practice, we recommend the biallelic estimator of F_ST_ in most cases (e.g. for assessing population structure), while the multiallelic-aware estimator of F_ST_ may be used as a descriptive measure of observed differentiation, or to standardize comparisons of sets of highly diverged populations. To facilitate this, we provide an extra flag --fst_biallelic, that can be paired with -- include_multiallelic_snps, to produce multiallelic-aware estimates of π and d_xy_ (which are not affected by homoplasy in the same way and are simply better estimators), but constrain F_ST_ to the biallelic estimator (when the coalescent F_ST_ is the estimand of interest, which should be most of the time).

### pixy in the contemporary toolkit

pixy combines a variety of features that make it useful across a wide variety of datasets: built-in missingness-aware statistics, support for arbitrary and variable ploidy, support for multiallelic sites, a full set of within- and between-population statistics (π, d_xy_, F_ST_, Watterson’s θ, Tajima’s D), all in one simple command-line workflow. We are not aware of another tool that currently offers this full suite of features. Table S4 compares pixy with related tools across input, missing-data handling, statistics, ploidy, and multiallelic support.

The closest contemporary to pixy currently is clam (Mirchandani et al. 2025), which estimates the denominators of π and d_xy_ from BAM files rather than from an all-sites VCF, and pairs this with a variants-only VCF to recover the numerators. The current version of pixy contains experimental support for this approach via a companion package, wisp (Samuk 2026). This method avoids the process of generating an unwieldy all-sites VCF, which can be computationally intensive. The approach of clam is attractive, and the estimates of pixy and clam agree under idealized simulated conditions (Mirchandani et al. 2025). A key issue with the approach of clam is that its callability is estimated using depth-based cutoffs of primary alignments, whereas variant sites are determined via a statistical variant calling process that often involves realignment and local reassembly of the underlying haplotypes (which may greatly change the canonical alignment in a particular region). This potentially creates divergence in statistical ascertainment between variant and invariant sites, the effects of which are unclear, particularly for haplotype-based variant callers. All-sites VCFs largely avoid this, by passing both variant and invariant sites through the same realignment and calling workflow. We note that this standardizes the calling process rather than every downstream annotation: variant-quality annotations are defined only for variant records, and so cannot be applied identically to both classes of site. Further work is needed to compare these approaches on empirical data, as existing comparisons rely on idealized simulations (Mirchandani et al. 2025).

### Limitations

There are several limitations of pixy worth noting. We note first what the phrase “unbiased with respect to missing data” does and does not claim. pixy conditions its denominators correctly on the site and genotype missingness represented in an all-sites VCF, which removes the bias that motivated the tool. It cannot correct missingness that is itself allele-dependent — reference bias, genotype-calling error, or callability that varies with haplotype divergence — because that information is not present in the VCF. First, and most importantly, a limitation pixy shares with any tool that consumes a VCF built against a single linear reference genome is reference bias: reads carrying non-reference alleles map more poorly, which depresses apparent diversity in a way no downstream arithmetic can recover (Brandt et al. 2015; Günther and Nettelblad 2019; Thorburn et al. 2023; Akopyan et al. 2025). Depending on the species being analyzed, this bias is plausibly larger than any of the biases we observed here. Graph-based references (Garrison et al. 2018) may alleviate this, and pixy can in principle consume VCFs derived from graph alignments as those workflows become standard.

Another limitation is that pixy, as an explicit design choice, is only focused on computation of summary statistics, and performs no filtering of sites or genotypes. The estimates produced by pixy are thus only as good as the upstream variant calling and filtering. The reason for this design choice is straightforward: the parameter space for what constitutes “correct” filtering is huge, and there are no universal heuristics that apply across all studies. We thus consider filtering out of the scope of pixy and place the responsibility for making biologically sound decisions about filtering on the analyst, who has the best knowledge of both the organism and the sampling design. There is an excellent literature on the importance of filtering in population genomics (e.g. Brault et al. 2026), and there are many existing tools to implement best practices (e.g. Danecek et al. 2021). Finally, pixy infers ploidy from the first records of each contig, and so assumes ploidy is constant within a contig and uniform across samples for that contig; datasets with within-contig ploidy transitions, such as PAR and non-PAR regions of a sex chromosome, must be split across contigs or files.

### Future directions

Along with continuing to maintain and update the software, three extensions to pixy are planned. The first is the computation of summary statistics directly from genotype likelihoods, which would extend pixy’s missingness-aware framework into the low-coverage regime currently served by ANGSD (Korneliussen et al. 2014). Secondly, we plan to provide estimates of uncertainty for all summary statistics via bootstrap confidence intervals or via a Bayesian approach (producing posterior distributions rather than point estimates). Finally, a key area of development for summary statistics is a unified framework for quantifying diversity in SNPs and structural variants, which is an active area of work.

## Conclusion

pixy broadens the original π and d_xy_ tool into a general toolbox for within- and between-population summary statistics in the presence of missing data, with support for arbitrary ploidy and multiallelic sites, multicore execution, a full test suite, and extensive documentation. We provide a validated, openly available implementation of the all-sites-VCF approach to estimating a full set of commonly used population genetic summary statistics that are unbiased with respect to missing data.

## Acknowledgments

We are very appreciative of the users and external contributors whose GitHub issue reports shaped this release, including the contributors of the missingness-aware estimators, the modularization and test infrastructure, and several correctness fixes. We especially thank Akira Hirao for the detailed identification of a subtle but important error in the computation of the standard deviation term used in our implementation of Tajima’s D. We thank the UC Riverside High-Performance Computing Center for maintaining the infrastructure and computational resources that facilitated our benchmarks.

AI Disclosure: Claude Code (Opus 4.7/4.8 and Sonnet 4.6) was used to annotate and reorganize the code and analysis pipelines for the statistical analyses and figures for this manuscript. This was done to ensure consistency and reproducibility for each subsection (authored by separate contributors). Starting in May 2026, Claude Code (Opus 4.7/4.8) was used to expand test coverage for pixy, optimize and document human-authored functions, and to format the conda-forge release recipes and dependency structure for python versions 3.12–3.13 using existing human-authored templates. GitHub Copilot (Haiku 4.5) was used to generate commit and pull request summaries for the software package pixy itself. All code and documentation altered by AI was manually checked and verified for accuracy by the lead author and passed through the human-authored continuous integration suite. The algorithms and original implementation of pixy are human-authored; AI assistance in the software was limited to the maintenance tasks described above. No AI was used in the writing of this manuscript, other than standard grammar checking tools and for assisting with bibliographic metadata checks, which were verified against source records.

## Funding

This work was supported by funding to K.S. (National Institutes of Health grant no. 5R35GM154837-03). G.L. was additionally supported by a National Science Foundation Graduate Research Fellowship Program Award.

## Data availability

pixy is available at https://github.com/ksamuk/pixy under the MIT license and is installable from conda-forge. Scripts used to perform the validation and analyses presented here are available at https://github.com/samuk-lab/pixy-2.0.0-analysis under the MIT license. Documentation is at https://pixy.readthedocs.io. The software itself is named pixy. “pixy 2” denotes the 2.x release line, and the exact release used for all analyses is 2.2.3. Work using the software should cite both Korunes and Samuk (2021) and the present paper.

Software archive. The exact pixy release used for the analyses here (2.2.3) is archived on Zenodo (DOI: 10.5281/zenodo.21385301); vcfsim (Goulart and Samuk 2026) is likewise archived (DOI: 10.5281/zenodo.21401147). A tagged release of the analysis repository is archived on Zenodo (DOI: 10.5281/zenodo.21634511).

Analysis code and workflows. All benchmark, simulation, and figure code is in the analyses/ and figures/ directories of the analysis repository (https://github.com/samuk-lab/pixy-2.0.0-analysis).

This includes the exact commands and workflow scripts for the simulations (vcfsim parameter tables and SLURM job scripts), the empirical VCF construction (GATK HaplotypeCaller through to all-sites VCFs), the pixy runs, and figure generation; the random seeds (and the per-arm seed-generation logic) are recorded alongside the simulation parameter tables. Per-figure summary tables are included in the repository, so all figures regenerate without rerunning the pipelines; large regeneratable intermediates (simulated VCFs, aggregated per-arm tables, and per-replicate Tajima’s D cells) are excluded but are rebuilt dynamically when running the analysis code.

Software versions. Analyses used pixy 2.2.3, with pixy 0.95.01 as the single-core legacy baseline in the multicore benchmark, and vcfsim (commit da0b8ad, two commits after v1.2.0) for the simulations, under Python 3.11–3.13. Tool versions for the empirical calling and filtering pipeline are given in the Empirical datasets methods above. Analysis-specific conda environments provide pinned versions of all software in the envs/*.yml files in each analysis module.

Benchmark hardware. Benchmarks were run on a single Intel Broadwell node of the UC Riverside HPCC (two Intel Xeon E5-2683 v4 CPUs @ 2.10 GHz, 32 physical cores, 450 GB RAM; SLURM intel partition). Benchmark jobs used 1–16 cores and 8 GB of RAM.

Empirical data. Empirical data were obtained from previously published sources. *Anopheles gambiae*: the Ag1000G project (*Anopheles gambiae* 1000 Genomes Consortium 2017), chromosome 3R mapped to the AgamP4 reference; the BFS and KES populations, 8 individuals each. *Arabidopsis arenosa*: the resequencing data of Monnahan et al. (2019) (BioProject PRJNA484107), mapped to the AARE701a reference (GCA_905216605.1; chromosome LR999451.1); the SPI and TRE populations, 8 individuals each. Sample accessions for all four populations are listed in Table S3.

## Supplementary material

**Table S1.** New features and major performance improvements in pixy, from the v0.95.02 release archived for Korunes & Samuk (2021) through pixy 2.2.3, by version, category, and feature.

| Version | Release date | Category | Feature | Description |
| --- | --- | --- | --- | --- |
| 1.0.0 | 2021-03-11 | Performance | NumPy vectorization | Core calculations vectorized with NumPy for $\approx 3\times$ single-core throughput. |
| 1.0.0 | 2021-03-11 | Performance | Native parallelization (`--n_cores`) | Parallelization implemented with the standard-library multiprocessing module. |
| 1.0.0 | 2021-03-11 | Performance | Lower, configurable memory (`--chunk_size`) | Memory greatly reduced; large windows chunked, small windows grouped. |
| 1.0.0 | 2021-03-11 | Windowing | BED file window support | Heterogeneous windows from a BED file. |
| 1.0.0 | 2021-03-11 | Windowing | Sites file support | Restrict computation to a tab-separated CHROM/POS sites list. |
| 1.0.0 | 2021-03-11 | Statistics | Site-level statistics (1 bp) | Per-site statistics at 1 bp resolution. |
| 1.0.0 | 2021-03-11 | Input | htslib/tabix as hard dependency | Requires bgzip-compressed, tabix-indexed VCFs for random access. |
| 1.0.0 | 2021-03-11 | Architecture | Zarr intermediate removed | Zarr intermediate genotype store removed for direct tabix access. |
| 1.0.0 | 2021-03-11 | Architecture | Genotype filtering removed | Filtration removed; users pre-filter with VCFtools/BCFtools. |
| 1.1.0 | 2021-05-10 | Statistics | Hudson's FST estimator (`--fst_type`) | Hudson (1992) FST selectable via `--fst_type` alongside Weir-Cockerham. |
| 1.2.1 | 2021-07-03 | Performance | Optimized per-site FST | Faster per-site FST; further small-window optimization in 1.2.2. |
| 1.2.5 | 2021-10-12 | Performance | Single-site mode optimization with sites file | Major `--window_size 1 + sites-file` speedups; explicit NAs in output. |
| 1.2.9 | 2024-02-06 | Input | GATK missing-data format support | $DP < 0$ GTs coerced to missing; variant scan widened to 100 kbp with a warning. |
| 2.0.0 | 2026-05-24 | Statistics | Unbiased Watterson's $\theta$ (--stats watterson_theta) | Missing-data-corrected estimator (Bailey et al. 2025). |
| 2.0.0 | 2026-05-24 | Statistics | Unbiased Tajima's D (--stats tajima_d) | Missing-data-aware estimator (Bailey et al. 2025); as of 2.0.0 the denominator is evaluated once from total $\eta$ and the mean per-site allele count (see New summary statistics). |
| 2.0.0 | 2026-05-24 | Statistics | Multiallelic site support (--include_multiallelic_snps) | Optionally include >2-allele sites; multiallelic FST handling fixed. |
| 2.0.0 | 2026-05-24 | Statistics | Arbitrary and variable ploidy | Any ploidy, variable across contigs; inferred from the first record of each contig. |
| 2.0.0 | 2026-05-24 | Input | CSI index support | Accepts both .tbi and .csi VCF indexes. |
| 2.0.0 | 2026-05-24 | Windowing | BED file 0-based half-open (breaking fix) | BED windows now 0-based half-open (breaking change from 1-based). |
| 2.0.0 | 2026-05-24 | Output | FST and Tajima's D component output | --fst_components and --tajima_components emit estimator components. |
| 2.0.0 | 2026-05-24 | Performance | $\approx 13\times$ faster cold start, $\approx 13\times$ lower startup memory | $\approx 13\times$ faster cold start and lower startup memory via deferred imports. |
| 2.0.0 | 2026-05-24 | Performance | $\approx 2.5\times$ faster overall; $\approx 27\%$ lower memory | $\approx 2.5\times$ faster, $\approx 27\%$ less memory via vectorized, pandas-free aggregation. |
| 2.0.0 | 2026-05-24 | Performance | Streamed aggregation (memory flat in window count) | Streaming aggregator keeps memory flat as window count grows. |
| 2.0.0 | 2026-05-24 | Performance | Batched per-contig ploidy inference | Per-contig ploidy inference batched into one tabix call. |
| 2.0.0 | 2026-05-24 | Dependencies | Dropped pandas, scipy, multiprocessing | pandas, scipy, and multiprocessing replaced with stdlib equivalents. |
| 2.0.0 | 2026-05-24 | Dependencies | Python 3.10–3.14 supported | Supported Python range broadened to 3.10–3.14. |
| 2.0.0 | 2026-05-24 | Security | Removed shell calls (command-injection surface) | Shell invocations replaced with pure-Python; removes injection surface. |
| 2.0.0 | 2026-05-24 | Packaging | Poetry build system + ruff + mypy + CI | Poetry build, ruff, mypy, and full CI on every PR. |
| 2.1.0 | 2026-06-03 | Input | wisp population-count mask support. | Reads per-population callable-site masks from wisp (experimental). |
| 2.1.2 | 2026-06-12 | Performance | Streaming FST-only reader | FST-only stream genotypes via htlib, lowering peak RSS. |
| 2.2.0 | 2026-06-16 | Input | GVCF support | Reads multi-sample GVCFs directly. |
| 2.2.2 | 2026-07-15 | Statistics | Corrected multiallelic Hudson's FST | Hudson's FST estimation fixed at sites with more than two alleles. |
| 2.2.2 | 2026-07-15 | Statistics | Biallelic FST with multiallelic $\pi$/dxy (--fst_biallelic) | Restrict FST to biallelic sites while $\pi$ and dxy use all sites, including multiallelic. |
| 2.2.2 | 2026-07-15 | Statistics | Mutation-based Watterson's $\theta$ and Tajima's D | $\theta_W$ and Tajima's D count mutations (Tajima's $s^*$), not segregating sites, at multiallelic sites; relatedly, alternate-fixed sites are no longer counted as segregating (affects the default biallelic path). |

**Table S2.** Accuracy of pixy’s estimators relative to their theoretical expectations, across the simulated ploidy × missingness grid of Figure 3. Hudson’s F_ST_ is the biallelic estimator, matched to its infinite-sites expectation; the other four statistics are estimated in multiallelic-aware mode. For each summary statistic, bias is the difference between the mean estimate and its theoretical expectation (“E”), pooled over the full grid of ploidies (2n, 4n, 6n, 8n) and missingness levels (0, 10, 25, 50, 75%; 72,000 simulation replicates per statistic). MCSE is the Monte Carlo standard error of the bias (SD/√n_sim) — the precision with which bias of the estimator is able to be measured given the number of replicate simulations (Morris et al. 2019). Relative bias expresses the bias as a percentage of E. Tajima’s D has E ≈ 0; its estimate and expectation are therefore shifted by +1 before the ratio is formed. Worst-cell bias is the largest deviation from the theoretical value in any single ploidy × missingness combination; for Tajima’s D this entry is an absolute deviation rather than a percentage.

| Statistic | n | Bias (mean — E) | Relative bias | MCSE | Worst-cell bias |
| --- | --- | --- | --- | --- | --- |
| $\pi$ | 72,000 | $-4.51 \times 10^{-6}$ | −0.013% | $3.0 \times 10^{-6}$ | 0.11% |
| $d_{xy}$ | 72,000 | $-4.12 \times 10^{-7}$ | −0.001% | $4.5 \times 10^{-6}$ | 0.05% |
| Hudson's $F_{ST}$ | 72,000 | $-1.27 \times 10^{-4}$ | −0.054% | $3.0 \times 10^{-5}$ | 0.15% |
| Watterson's $\theta_W$ | 72,000 | $-3.07 \times 10^{-6}$ | −0.008% | $2.6 \times 10^{-6}$ | 0.07% |
| Tajima's D | 72,000 | $-1.72 \times 10^{-4}$ | −0.018% | $5.5 \times 10^{-4}$ | 0.0028 |

**Table S3.**
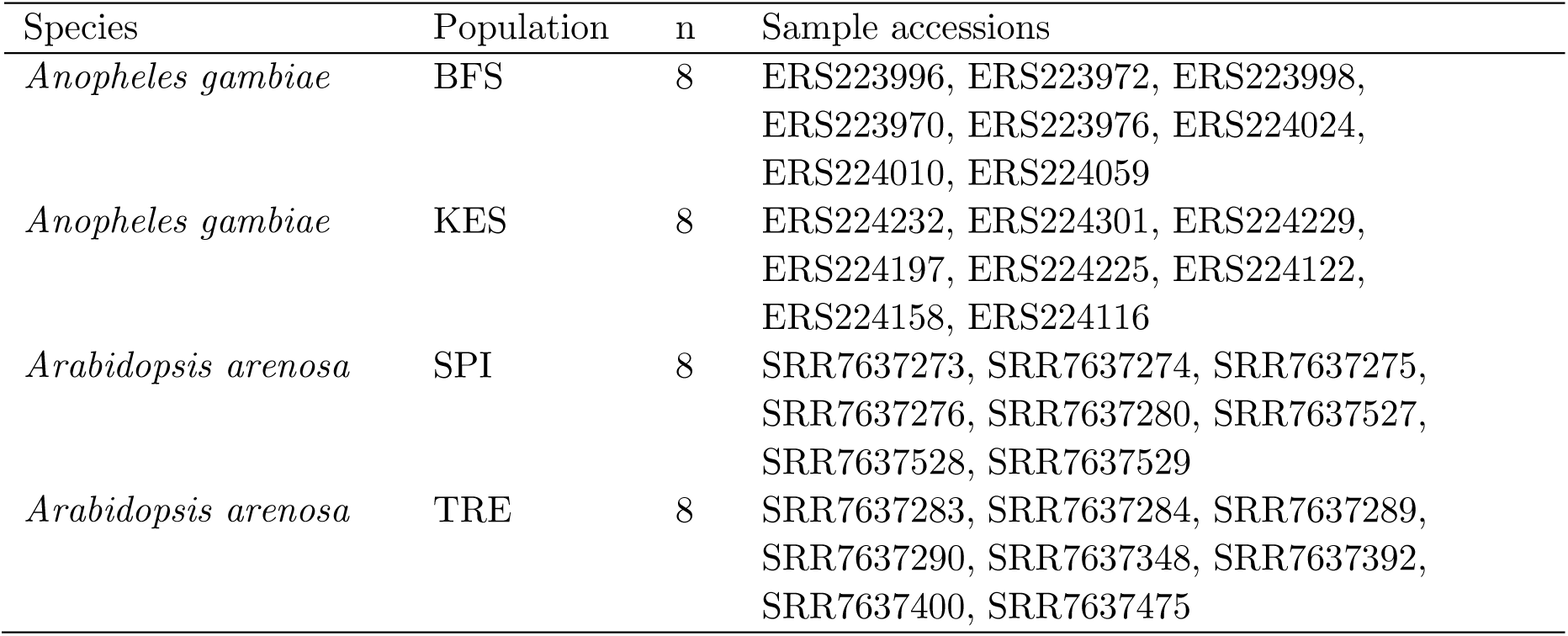
Sample accessions for the empirical analyses. *Anopheles gambiae* samples are ENA sample accessions from the Ag1000G project (chromosome 3R, AgamP4 reference); *Arabidopsis arenosa* samples are SRA run accessions from BioProject PRJNA484107 (Monnahan et al. 2019; AARE701a reference, GCA_905216605.1, chromosome LR999451.1).

**Table S4.**
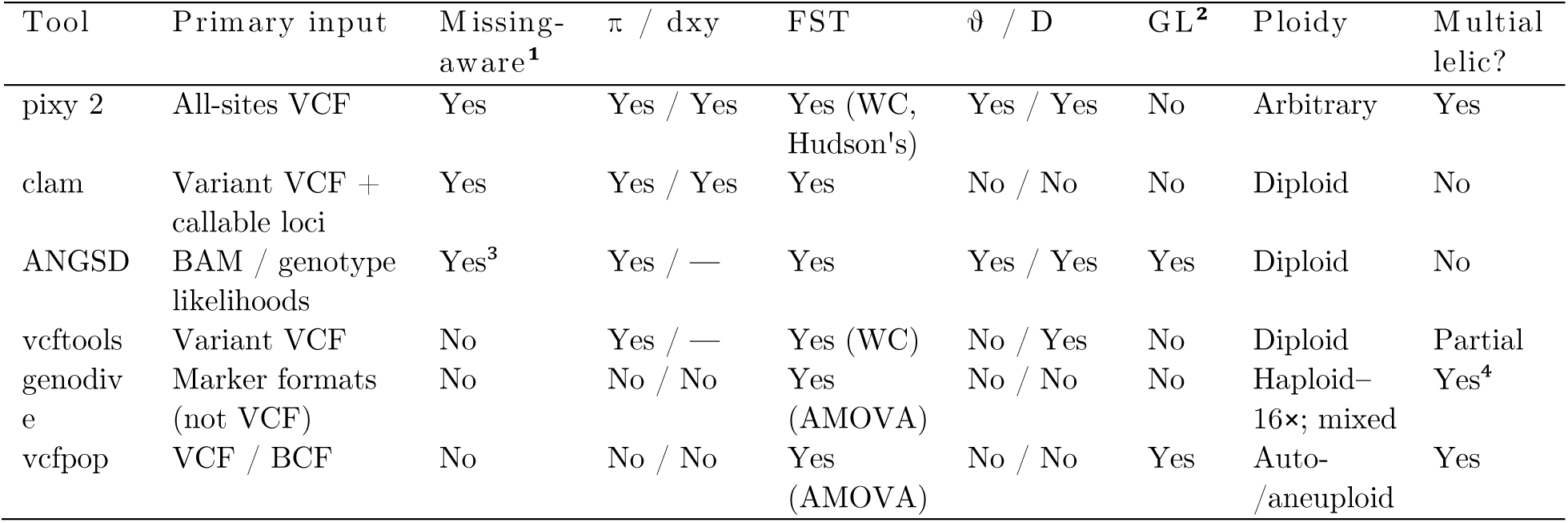
Feature comparison of pixy and related population-genomic tools. ¹“Missing-aware” = Correct denominators via all-sites or callable-loci information. ²GL: operates on genotype likelihoods (low-coverage-aware). ³ANGSD works from read data or likelihoods rather than called genotypes, treating missing data as genotype uncertainty. ⁴genodive handles multiallelic markers but is not a genome-scan tool. Windowed genome scans are supported by pixy (fixed, BED, sites-file), clam, ANGSD (sliding), and vcftools (fixed); genodive and vcfpop are marker/structure-oriented and do not scan windows. Interfaces are command-line except genodive (macOS GUI). Compiled from each tool’s primary publication: pixy (Korunes and Samuk 2021), clam (Mirchandani et al. 2025), ANGSD (Korneliussen et al. 2014), vcftools (Danecek et al. 2011), genodive (Meirmans 2020), vcfpop (Huang et al. 2025).

**Figure S1.**
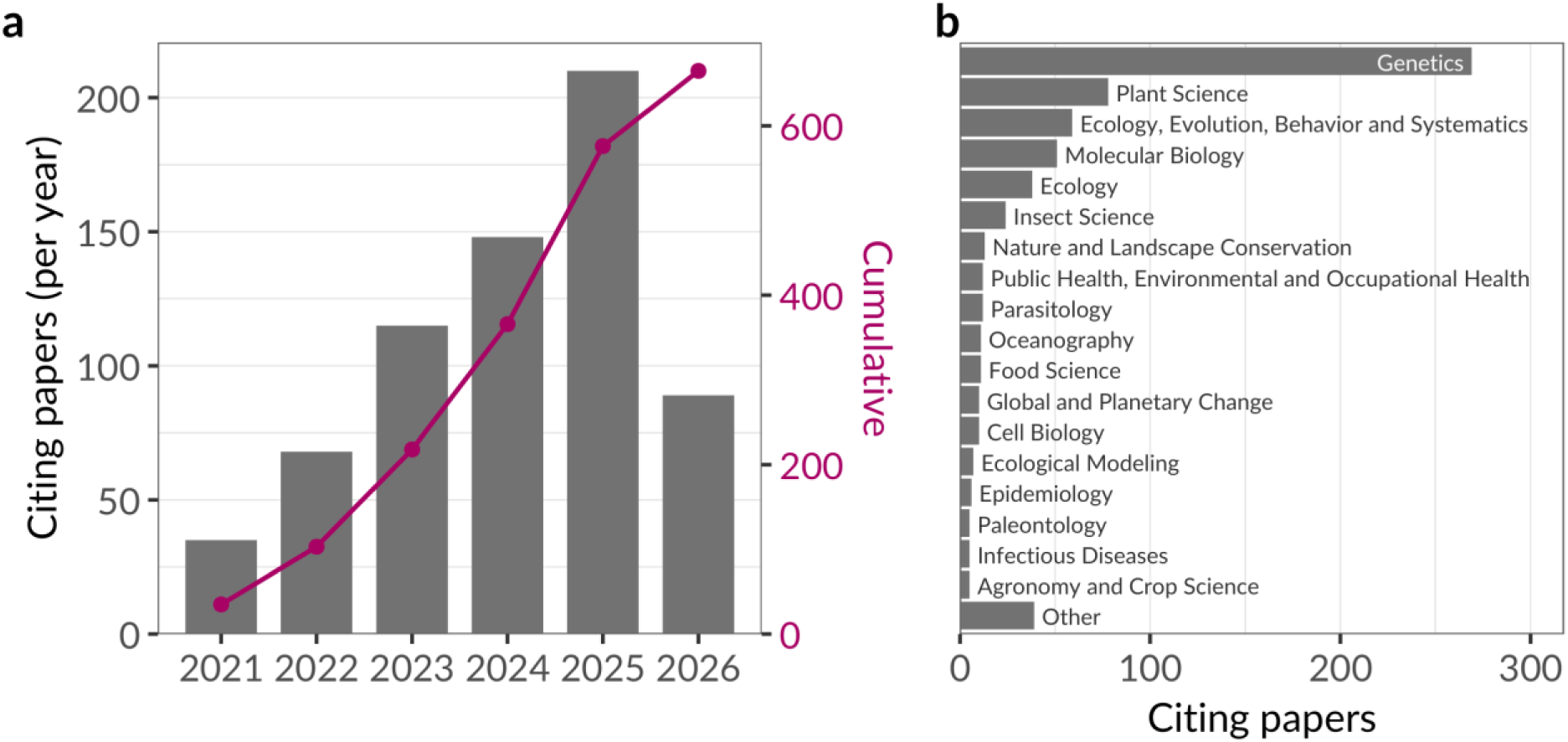
Citation footprint of pixy. (a) Citing papers per year (bars) with cumulative total (line). (b) Distribution of citing papers across OpenAlex subfields; subfields with fewer than five citing papers are pooled into “Other”.

**Figure S2.**
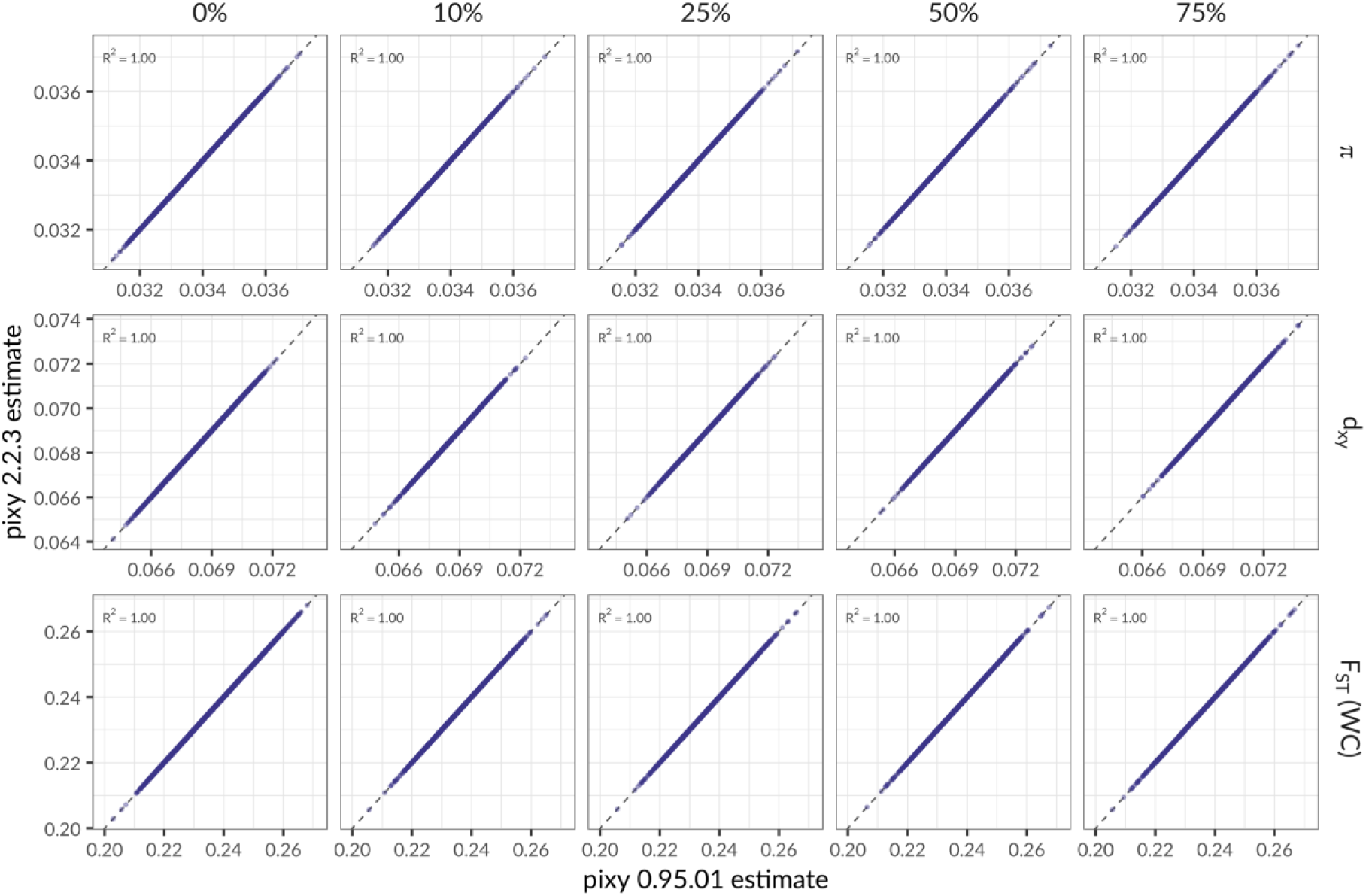
Diploid cross-version agreement. Per-VCF estimates from pixy 0.95.01 (x) versus pixy 2.2.3 (y) for π, d_xy_, and Weir–Cockerham F_ST_ (rows) across missingness 0–75% (columns); each point is one simulated replicate, dashed line is 1:1.

## Supplementary note: Finite-sites expectations for Watterson’s **θ** and Tajima’s D under multiallelic, polyploid sampling

As of version 2.0.0, pixy reports the missingness-aware Watterson’s θ and Tajima’s D of Bailey et al. (2025), with minor modifications for Tajima’s D (see Methods section). Here we describe one issue specific to multiallelic, polyploid sampling, and how the finite-sites theoretical expectations shown in Figure 3 were determined, which ended up being nontrivial.

### The problem

In multiallelic-aware mode pixy estimates π from pairwise differences between genotypes but, by default, estimates Watterson’s θ from the number of segregating sites. Under an infinite sites (and neutral) model these are the same quantity, but under a finite sites model they are not. A site with more than one mutation in its genealogy can segregate for three or four alleles (increasingly likely as more gene copies are sampled) so the site count S and the mutation count η diverge:

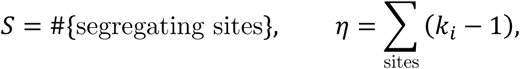

where kᵢ is the number of alleles at site i and η, the parsimony-minimum number of mutations, is Tajima’s (1996) s*. The multiallelic-aware π estimator implemented in pixy counts these extra mutations while a site-based θ does not, so π exceeds Watterson’s θ and the Tajima’s D that contrasts them is biased upward, increasingly so with ploidy (the positive, ploidy-increasing D of the stock statistic across the polyploid arms, ≈ +0.07 to +0.18). Following Tajima (1996) and Bailey et al. (2025), the fix is to substitute η for S in both the numerator and the denominator of D (the denominator substitution follows Misawa and Tajima 1997), evaluated at the gene copies actually observed at each site so that missing data and mixed ploidy are handled correctly. On biallelic data this reduces exactly to the classical Tajima (1989) statistic. The correction is consistent rather than unbiased per se because η ≥ S, and thus it can only shift D downward. D thus settles on a small finite-sites floor rather than zero, which is an intrinsic residual with no general closed form (Roychoudhury and Wakeley 2010; Bhaskar, Kamm and Song 2012).

### Deriving a theoretical expectation under finite sites and missing data

The θ and π theoretical reference lines in Figure 3 are the finite-sites JC69 expectations of Tajima (1996). These are the values against which pixy’s estimates are compared; they are not quantities that pixy itself computes. In multiallelic mode pixy estimates π exactly as it does in biallelic mode, from pairwise differences among the observed genotypes, and multiallelic sites simply contribute their additional pairwise comparisons. For nucleotide diversity,

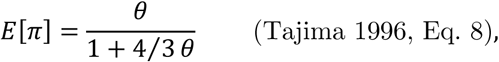

and each Watterson expectation is the corresponding expected mutation count divided by the harmonic number a₁(n):

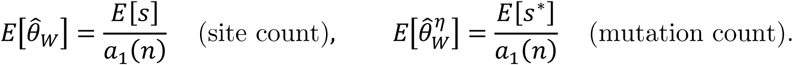

where E[s] and E[s*] are the JC69 closed forms of Tajima (1996, Eqs. 6 and 15). Because s* recovers only the minimum mutations implied by the observed alleles, the mutation-count expectation is ≈ 4% below the true θ = 4Neμ, an irreducible homoplasy floor. In our simulations, we reproduced every entry of Tajima (1996)’s Table 1 to four decimal places, and the simulated means match to within 4 × 10⁻⁵ across ploidy. Evaluating a₁(n) at the per-site copy number actually observed, rather than the nominal ploidy × sample size, makes the θ reference slope upward with missingness, and this was recovered analytically with no re-simulation.

To reiterate: Tajima’s D itself has *no convenient finite-sites closed form*, and its denominator is random and mutation-model-dependent (Tajima 1989). The single-locus analytic variance does not apply here because the simulations recombine within windows (like actual biological windows). We therefore set the D reference by simulation, using the mean of the corrected statistic over a neutral coalescent null matched to each cell’s window length, recombination rate, ploidy, sample size, and missingness (msprime with a JC69 discrete genome). This null model’s mutation rate was fixed by inverting the π expectation above, 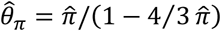, so that the simulated θ matches the data. This absorbs both the bias and the variance of the contrast under finite sites. The resulting expected Tajima’s D has a small negative floor (−0.033 at 2n, shallowing to −0.016 at 8n) on which the corrected estimates sit, whereas the previously implemented site-counting statistic rises with ploidy, a signature of the counting mismatch the correction removes.

The full scripts and tools used to derive these expectations are available along with the rest of the analysis code here: https://github.com/samuk-lab/pixy-2.0.0-analysis.

## Notes

https://github.com/ksamuk/pixy

### Competing Interest Statement

The authors have declared no competing interest.

